# Single-cell immune repertoire sequencing of B and T cells in murine models of infection and autoimmunity

**DOI:** 10.1101/2022.02.07.479381

**Authors:** Danielle Shlesinger, Kai-Lin Hong, Ghazal Shammas, Nicolas Page, Ioana Sandu, Andreas Agrafiotis, Victor Kreiner, Nicolas Fonta, Ilena Vincenti, Ingrid Wagner, Margot Piccinno, Alexandre Mariotte, Bogna Klimek, Raphael Dizerens, Marcos Manero-Carranza, Raphael Kuhn, Roy Ehling, Lester Frei, Keywan Khodaverdi, Camilla Panetti, Nicole Joller, Annette Oxenius, Doron Merkler, Sai T. Reddy, Alexander Yermanos

## Abstract

Adaptive immune repertoires are composed by the ensemble of B and T cell receptors (BCR, TCR) within an individual and reflect both past and current immune responses. Recent advances in single-cell sequencing enable recovery of the complete adaptive immune receptor sequences in addition to transcriptional information. Such high-dimensional datasets enable the molecular quantification of clonal selection of B and T cells across a wide variety of conditions such as infection and disease. Due to costs, time required for the analysis and current practices of academic publishing, small-scale sequencing studies are often not made publicly available, despite having informative potential to elucidate immunological principles and guide future-studies. Here, we performed single-cell sequencing of B and T cells to profile clonal selection across murine models of viral infection and autoimmune disease. Specifically, we recovered transcriptome and immune repertoire information for polyclonal T follicular helper cells following acute and chronic viral infection, CD8+ T cells with binding specificity restricted to two distinct peptides of lymphocytic choriomeningitis virus, and B and T cells isolated from the nervous system in the context of experimental autoimmune encephalomyelitis. We could relate repertoire features such as clonal expansion, germline gene usage, and clonal convergence to cell phenotypes spanning activation, memory, naive, antibody secretion, T cell inflation, and regulation. Together, this dataset provides a resource for experimental and computational immunologists that can be integrated with future single-cell immune repertoire and transcriptome sequencing datasets.

## Introduction

B and T cells play a central role in orchestrating the immune response by recognizing foreign antigens through their B cell receptor (BCR; secreted version: antibodies) and T cell receptor (TCR), respectively. Upon primary antigen exposure, B and T cells rapidly undergo clonal expansion and can adopt a diverse set of cellular phenotypes with corresponding effector functions (Lu et al. 2017; Sandu et al. 2020; Utzschneider et al. 2016). B cells further engage in germinal center (GC) reactions, where they undergo diversification through class-switch recombination and somatic hypermutation, hence increasing the range of effector functions and molecular recognition (Li et al. 2004; Georgiou et al. 2014). B cells expressing high-affinity BCRs are selected for by T follicular helper (Tfh) cells, which are characterized by their expression of CD4, PD-1, CXCR5, and BCL6, and thus provide the required selection signals for B cell differentiation (Klein and Dalla-Favera 2008; Victora and Nussenzweig 2012; Mesin, Ersching, and Victora 2016; Stebegg et al. 2018; Vinuesa et al. 2016; Crotty 2011; Tangye et al. 2013; Biram, Davidzohn, and Shulman 2019). Following the resolution of immune challenges, and thereby the diminishing presence of cognate antigens, B and T cells contract and preferentially adopt heterogenous memory phenotypes characterized by expression of certain gene signatures (Kalia, Sarkar, and Ahmed 2010; Welten et al. 2020). While this selection and differentiation of lymphocytes is heavily regulated under healthy conditions, dysfunctional clonal selection can contribute to autoimmunity and host pathology (Brandão et al. 2020; Constantinescu et al. 2011).

The development of standardized protocols, reagents and dedicated commercial platforms (e.g., 10X Genomics) has made it possible to perform single-cell immune repertoire sequencing that simultaneously recovers gene expression (whole transcriptome) and immune repertoire (full-length, paired chains of BCRs [heavy and light chains] and TCRs [alpha and beta chains]) information (Yermanos, Neumeier, et al. 2021; Yermanos, Agrafiotis, et al. 2021; Horns, Dekker, and Quake 2020; Khatun et al. 2021; Croote et al. 2018; Saikia et al. 2019; Singh et al. 2019; Neumeier et al. 2021; Agrafiotis et al. 2021; Zheng et al. 2017). This now enables investigation of how parameters such as clonal expansion relate to gene expression phenotypes (Csepregi et al. 2020), thereby providing novel insights not accessible by bulk sequencing (e.g., BCR variable heavy chain, TCR variable beta chain) methodologies (Welten et al. 2020; Yermanos et al. 2020; Chang et al. 2020). In recent years, a number of studies have employed single-cell immune repertoire sequencing to dissect the heterogeneity and differentiation of T and B cells in contexts such as infection, disease, and vaccination (Neumeier et al. 2021; Yermanos, Neumeier, et al. 2021; Sureshchandra et al. 2021; Mogilenko et al. 2021; Hiltensperger et al. 2021).

Practical considerations such as the necessary cost and time required to generate and analyze each single-cell sequencing sample has been a limitation to performing large-scale experiments (i.e., with many samples). These challenges combined with current practices of academic publishing may result in the loss of publicly available data, either due to low sample numbers as a result of practical or technical difficulties. Despite this, pilot experiments have the potential to nevertheless provide insight and be used as a comparison to future experiments, however, many of these datasets have not reached the public.

Here, we combine single-cell sequencing of both T and B cells from murine models of infection and autoimmune disease, exploring both gene expression and immune repertoire profiles. This dataset includes T follicular helper (Tfh) cells following acute and chronic infections with the model virus lymphocytic choriomeningitis virus (LCMV), CD8+ T cells with antigen-restricted specificity arising from spleens 15 months post infection with LCMV or murine cytomegalovirus (MCMV), thereby revealing distinct phenotypes of memory and inflationary T cells. These virus-specific T cells could be further compared to CD8+ T cells arising from the brain in a murine model of T-cell-driven neurological disease (Di Liberto et al. 2018), which revealed gene expression patterns resembling T cell exhaustion. Finally, we uncovered minimal transcriptional differences in a pilot study investigating the differences in BCR and TCR repertoires following induction of experimental autoimmune encephalomyelitis (EAE) using either myelin-oligodendrocyte glycoprotein (MOG) peptide (MOG35-55) or recombinant protein (rMOG). Our collection of single-cell immune repertoire sequencing experiments not only supports previously published results but also uncovers novel principles of B and T cell clonal selection. Therefore, our dataset can serve as a reference for experimental and computational immunologists alike to inform future studies.

## Results

### Polyclonal Tfh cell populations following acute and chronic LCMV infection are transcriptionally heterogeneous, highly-expanded, and share clones with GP66-specific CD4+ T cells

Following immune challenges, B cells engage in germinal center (GC) reactions, where they undergo diversification through somatic hypermutation (SHM). B cells expressing high-affinity BCRs are selected for by Tfh cells and are characterized by their expression of CD4, PD-1, CXCR5, and BCL6 (Klein and Dalla-Favera 2008; Victora and Nussenzweig 2012; Mesin, Ersching, and Victora 2016; Stebegg et al. 2018; Vinuesa et al. 2016). In an acute LCMV infection, viral clearance occurs within 8-10 days and is dependent on activation of cytotoxic CD8+ T cells (Vella, Herati, and Wherry 2017; Matloubian, Concepcion, and Ahmed 1994). However, during a chronic LCMV infection, T cell exhaustion occurs, making clearance dependent on late-emerging neutralizing antibodies that appear months after infection (Ciurea et al. 2000; Eschli et al. 2007; Battegay et al. 1993). Tfh cells have been shown to be key players in the production of these late-emerging neutralizing antibodies and the resolution of chronic infection (Fahey et al. 2011; Greczmiel et al. 2017). To explore transcriptional and repertoire heterogeneity of Tfh cells during acute and chronic LCMV infection, we isolated CD4+CXCR5+PD-1+ T cells and performed single-cell sequencing of the TCR repertoire and transcriptome (Figure 1A). Cells were isolated by magnetic-activated cell sorting (MACS) from the spleens of C57BL/6 mice infected with LCMV causing acute (200 ffu LCMV clone 13 i.v., 8 dpi n=2) or chronic disease (2×10^6^ ffu LCMV clone 13 i.v., 8, 25 and 50 dpi, n=2 at each time point) (Kräutler et al. 2020). Following alignment to reference murine transcriptome, we recovered gene expression data (GEX) for 55,440 cells with an average of 1,186 median genes per cell (Figure S1A).

**Figure 1.**
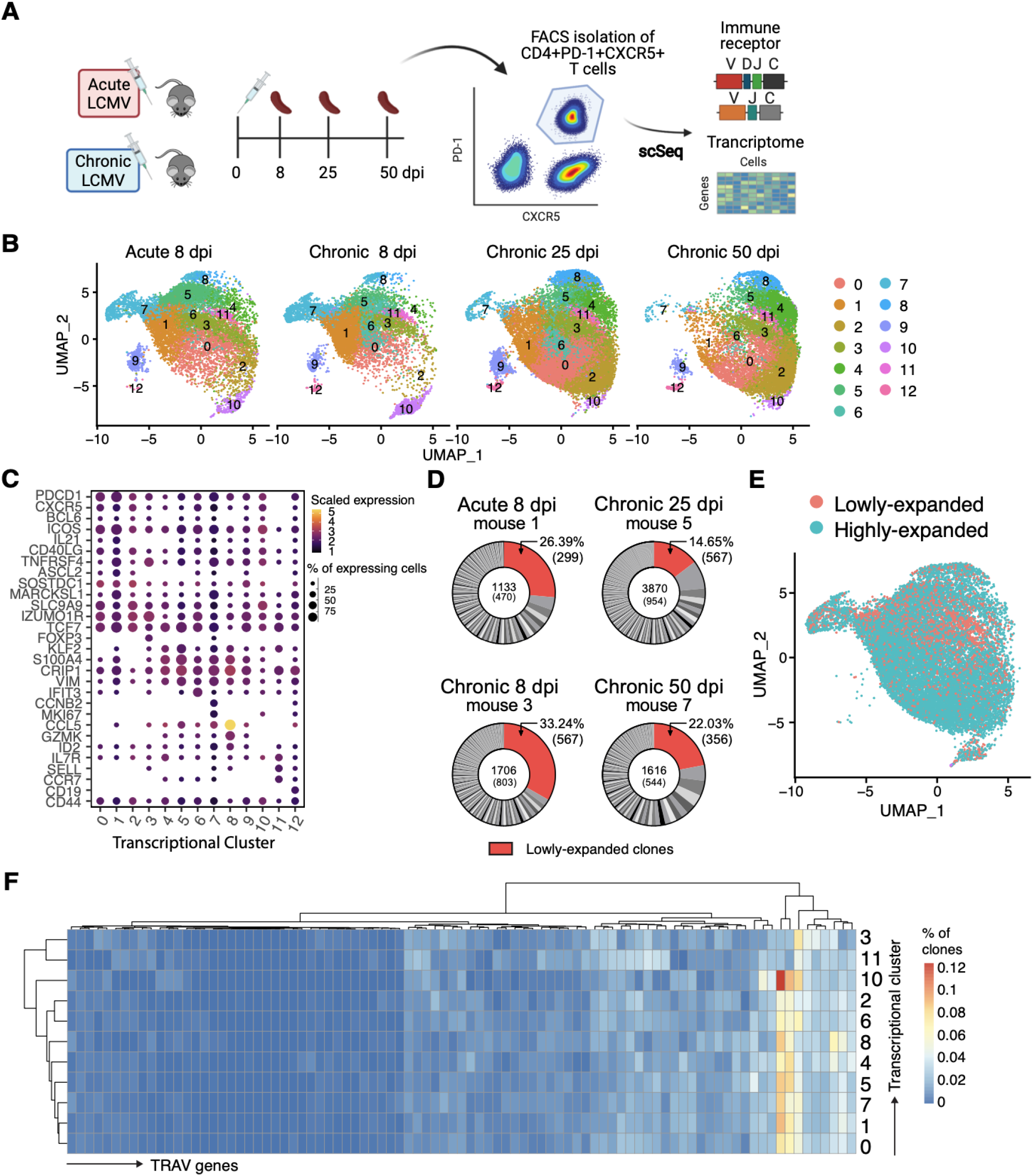
Different transcriptional phenotypes emerge in polyclonal Tfh cell populations following acute and chronic LCMV infection. Experimental Setup. CD4+CXCR5+PD-1+ cells were Isolated from spleens of C57BL/6 mice infected with acute LCMV (n=2) or chronic LCMV infection (n=6). Each time point included two biological replicates. Cells were then sorted and sequenced to obtain both transcriptomes and TCR repertoires. B. Uniform manifold approximation projection (UMAP) separated by experimental group. C. Dottile plot depicting scaled expression of genes of interest separated by transcriptional clusters. D. Representative donut plots depicting distribution of clonal expansion in each infection time point. Each section corresponds to a unique clone (defined by CDR3α-CDR3β nucleotide (nt) sequence) and the size corresponds to the fraction of cells relative to the total repertoire. Lowly-expanded clones (supported by only one unique cell) are colored in red. E. UMAP displaying clonal expansion of Tfh cells after acute and chronic infection. Clones are defined by CDR3α-CDR3β nt sequence. Lowly-expanded clones are represented by only one unique cell, while highly-expanded clones are supported by more than one cell. F. Heatmap depicting percentage of clones (defined by CDR3α-CDR3β nt sequence) using a particular TRAV gene (rows) in each cluster (columns).

We next used uniform manifold approximation projection (UMAP) to group cells with similar gene expression profiles into clusters (Figure 1B), thus revealing a variety of transcriptional phenotypes such as “Tfh-like” (*Sostdc1, Marcksl1,Slc9a9*), T follicular helper (Tfh) and regulatory cells (Tfr; *Foxp3, Bcl6*), Th1- or T central memory precursor (Tcmp) like (*Crip1, Klf2, S100a4, Vim*), interferon-stimulated gene expressing (*Ifit3, Isg15*), proliferative (*Ccnb3, Mki67*), Th1-like (*Ccl5, Nkg7, Id2, Gzmk*), PI3K pathway expressing and naive-like (*Sell, Ccr7, Il7r* and low *Cd44*) (Figure 1C, Figure S1B, C). We additionally detected two clusters expressing genes associated with B cells and the complement cascade, suggesting minor contamination (clusters 9 and 12) (Figure S1B). Cells belonging to mice sacrificed at 8 dpi with either acute or chronic LCMV (early time points of infection) occupied a distinct transcriptional space compared to cells belonging to mice with chronic LCMV sacrificed at 25 or 50 dpi (later time points of infection) (Figure S2A). This was characterized by increased expression of genes associated with fatty acid metabolism, hypoxia and *Myc* targets, suggesting a more activated and proliferative phenotype early after infection (Figure S2B). This is consistent with previous reports of Tfh cells proliferating two weeks after protein immunization (Merkenschlager et al. 2021).

We next aimed to explore the clonal diversity of Tfh cells during acute or chronic LCMV infection. For this, only cells expressing a single TCR alpha (α) and beta (β) chain were retained in the analysis and cells belonging to those clusters likely arising from contamination (9 and 12) were similarly excluded. We observed a polyclonal population of Tfh cells and an overall high degree of expansion in both acute and chronic LCMV infection (Figure 1D, S2C). Notably, expansion was highest at 25 dpi with chronic LCMV and decreased again at 50 dpi, potentially representing T cell contraction occurring as the infection is resolved (Kräutler et al. 2020). We further analyzed the transcriptional space occupied by lowly- and highly-expanded clones (represented by one unique cell or more than one cell respectively), which revealed that Tfr cells and naïve cells especially were lowly-expanded, although all clusters contained some lowly-expanded cells (Figure 1E, S2D).

The abundance of certain TCR V germline genes or their pairings have been previously observed in virus-specific T cells in several infection conditions (Qu et al. 2016; Izraelson et al. 2018; Welten et al. 2020; Yermanos et al. 2020; Khatun et al. 2021; Kuhn et al. 2022). Additionally, a previous study reported preferential usage of TRAV-TRAJ gene combination in Th1 and Tfh cells, indicating a contribution of the TCR α chain to lineage commitment (Khatun et al. 2021). This is congruent with previous findings demonstrating that Tregs have been reported to have distinct germline gene usage (e.g. dominated by TRBV5) and that Tfr have different TCR specificities than Tfh (Maceiras et al. 2017; Schorer et al. 2020). Here, clusters 3 (Tfr) and 11 (naive-like) could be distinguished from other clusters by hierarchical clustering according to their TRAV gene usage (Figure 1F), suggesting a difference in T cell lineages compared to the other clusters. On an individual-chain level, similar TCR β V genes were expressed across all infection conditions (Figure S3A, S3B), although on a clonal level, no group-specific pattern of TRAV-TRBV pairings could be observed (Figure S3C). This is in contrast to previously reported results where TCR repertoires against a single peptide demonstrated stereotypic germline gene usage (Kuhn et al. 2022), suggesting that polyclonal repertoires encoding diverse TCR specificities are less dominated by certain combinations of germline genes.

To investigate clonal convergence we expanded our analysis by including previously published TCR repertoires of protein immunized mice and mice infected with a different LCMV strain. To this end, we integrated CD4+CD44+GP66+ T cells from mice infected with acute LCMV (Armstrong strain, 10 dpi) (Khatun et al. 2021) and CD4+CD62^low^CD44^high^CXCR5^high^PD-1^high^BCL6+ Tfh cells from hemi-splenectomized mice following immunization with ovalbumin (OVA) (7 and 21 days post immunization) (Merkenschlager et al. 2021) (Figure 2A). Hierarchical clustering according to TRAV gene usage separated the different data sets (Figure 2B). Notably, Tfh cells from both virus infection and OVA immunization clustered in a separate node than CD4+CD44+GP66+ T cells, again in line with previous reports relating the TCR α chain to CD4+ cell fate decisions (Khatun et al. 2021). We then further explored repertoire similarities across mice in the context of shared clones after defining clones by identical CDR3α and CDR3β amino acid (aa) sequences. Mice infected with LCMV clone 13 (acute and chronic) shared up to 20 clones with each other and at least one clone from the GP66+ repertoires. Surprisingly, mice with chronic LCMV 8 and 25 dpi shared 6 clones with OVA immunized mice (Figure 2C). Visualizing all public clones on the UMAP showed that public clones were mainly absent from the Tfr and naive-like clusters (clusters 3 and 11) (Figure 2D, S3D). As Tfr cells have been suggested to have autoreactive TCR specificities (Maceiras et al. 2017), our data suggest that Tfr clones might therefore also show a more personalized response.

**Figure 2.**
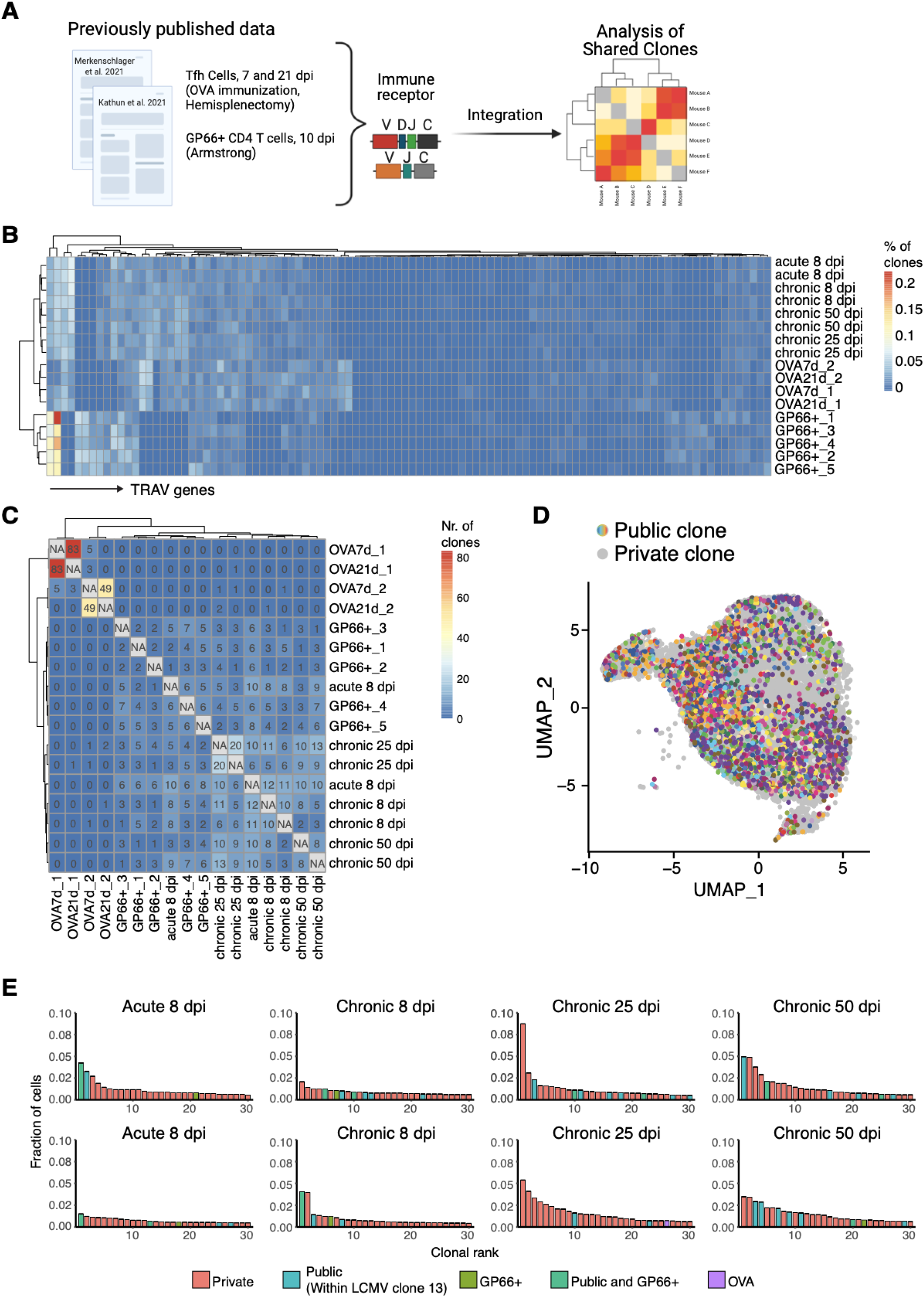
Polyclonal Tfh cells following acute and chronic infection include highly-expanded, public GPC-specific clones. **A**. Graphical illustration of the integration of previously published data in order to investigate public clones: GP66-specific CD4+ cells from mice infected with Armstrong and sacrificed 8 dpi (Khatun et al. 2021) and Tfh cells from OVA-immunized, hemi-splenectomized, mice (7 and 21 days post immunization) (Merkenschlager et al. 2021). B.Heatmap depicting percentage of clones (defined by CDR3α-CDR3β nt sequence) using a particular TRAV gene (rows) in each sample (columns). C. Number of identical clones (defined by CDR3α-CDR3β nt sequence) found between Tfh cells from mice infected with LCMV clone 13, mice infected with Armstrong sacrificed 8 dpi (Khatun et al. 2021) and Tfh cells from OVA-immunized, hemi-splenectomized, mice (7 and 21 days post immunization) (Merkenschlager et al. 2021). D. Uniform manifold approximation projection (UMAP) displaying shared clones between between Tfh cells from mice infected with LCMV clone 13, mice infected with Armstrong sacrificed 8 dpi (Khatun et al. 2021) and Tfh cells from OVA-immunized, hemi-splenectomized, mice (7 and 21 days post immunization) (Merkenschlager et al. 2021). E.Top 30 most expanded clones (defined by CDR3α-CDR3β nt sequence). Private clones are colored in red. Clones shared between mice infected with clone 13 (acute or chronic) are colored in blue and termed “Public”. Clones colored in green have inferred GP66 specificity. Clones colored in teal are “Public” and have inferred GP66 specificity. Purple colored clones are shared with OVA-immunized, hemi-splenectomized, mice (7 and 21 days post immunization) (Merkenschlager et al. 2021).

We next questioned whether these repertoire similarities across mice were also observed in the context of public clones. We further expanded our analysis of clonal convergence by including previously published TCR repertoires of protein immunized mice or mice infected with a different LCMV strain. Next, we investigated the extent of clonal expansion of all shared clones by looking for public clones within the top 30 most expanded clones. Taking advantage of the known epitope specificity of GP66+ T cell repertoires, we specifically highlighted clones shared with the GP66+ T cell repertoires to infer TCR specificities. Several highly expanded clones were public across all mice infected with LCMV clone 13 (Figure 2E). Although we did not sort for virus-specific cells, the expansion of some virus-specific Tfh cells could nevertheless be confirmed, as several highly-expanded clones shared an identical CDR3α-CDR3β aa sequence with GP66+ T cells. Interestingly, one clone from the OVA immunized mice was present in the top 50 most expanded clones of one repertoire arising 25 dpi with chronic LCMV (Figure 2E). This particular clone, however, was represented by a single cell in one of the OVA immunized mice, which may arise due to the previously described bystander-activation that has been suggested to lead to antigen-unspecific Tfh cells in OVA-immunized mice (Ritvo et al. 2018). Moreover, clonally expanded plasma cells with dual specificity to OVA and LCMV have previously been observed in chronically infected mice (Neumeier et al. 2021). Together, this raises the question whether this particular clone is indeed specific to both LCMV or OVA and further motivates the exploration of TCR specificities of Tfh cells arising following immune challenges.

### Memory, inflationary, and exhaustion signatures of virus-specific CD8+ T cells

We recently demonstrated that gp_33-41_-specific (GP33) CD8+ T cells adopted distinct transcriptional and selection phenotypes 28 days post acute, chronic, and latent viral infection (Kuhn et al. 2022). We questioned whether infection-specific repertoire fingerprints remained stable 15 months post infection (mpi). We therefore performed single-cell immune repertoire sequencing on splenic GP33+CD8+ T cells from age-matched mice infected with either LCMV Armstrong or two distinct murine cytomegalovirus containing the LCMV gp_33-41_ peptide in either the *m45* or *ie2* loci(MCMV-*m45*-gp33 and MCMV-*ie2*-gp33, respectively) (Figure 3A). Although both MCMV strains induce latent viral infections characterized by reactivation events, the location of gp_33-41_ allows for the isolation of CD8+ GP33-specific T cells that can be characterized by inflationary (using MCMV-*ie2*-gp33) and non-inflationary (using MCMV-*m45*-gp33) T cell phenotypes (Welten et al. 2020). This provided an age-matched comparison of virus-specific CD8+ T cells either lacking antigen stimulation versus repeated exposure to their cognate antigen. Overall, we recovered GEX data for 24,381 cells with an average of 956 median genes per cell (Figure 3A, S4A). Additionally, we compared these cell phenotypes to T cells arising from the viral déjà vu model, which has previously been leveraged to induce T-cell driven CNS inflammation (Di Liberto et al. 2018; Merkler et al. 2006; Kreutzfeldt et al. 2013). In this model, neonatal intracranial (i.c.) infection of an attenuated strain of LCMV (rLCMV) causes CNS-restricted viral persistence which leads to the development of immune-driven CNS disease following an i.v. infection with wild type LCMV (wtLCMV). We performed single cell immune repertoire sequencing on immunodominant NP396+ CD8+ T cells of mice receiving only a neonatal i.c. infection (carrier-non-challenge [CNC], n=1) and mice receiving both an i.c. infection and an i.v. infection with wtLCMV following 6 weeks and sacrificed after either 10 or 25 days (Carrier challenge [CC10 and CC25, respectively], n=3 for each time point) (Figure 3A). Here, we recovered GEX data for 36,590 cells with an average of 1,001 median genes per cell (Figure S4A). We then performed unsupervised clustering and UMAP on cells from both experiments in addition to our recently published GP33-specific CD8+ T cells 28 dpi (Kuhn et al. 2022). This resulted in 10 clusters, of which all with the exception of cluster 10 were represented by cells from all three experiments (Figure 3B, 3C). We next investigated the cluster-defining genes in an unbiased manner, thus observing clusters enriched in genes associated with phenotypes such as effector and inflationary (*Klrg1, Gzma, Zeb2)*, memory (*Il7r, Sell, Bcl2*), exhaustion (*Pdcd1, Ctla4, Tigit, Stat3, Tox*), interferon stimulated gene expressing (*Isg15, Ifit3*) and proliferation (*Pclaf, Mki67, Ccnb2*) (Figure 3D, S4B). Even 15 mpi, cluster 0 expressing genes associated with the previously reported effector-memory phenotype of inflationary CD8+ T cells could be clearly observed (Welten et al. 2020) (Figure 3C, 3B, 3D, S4B). Memory clusters 1 and 5 were represented by a higher proportion of cells at 15 mpi and CC25 compared to their earlier infection counterparts. Notably, chronic LCMV infection 28 dpi as well as CC10 and CC25 had a higher fraction of cells belonging to cluster 2 and 7, expressing genes associated with exhaustion and proliferation, respectively. Comparing age-matched mice within cluster 0 (effector and inflationary) and cluster 1 (memory), revealed *Cdk8, Lars2, Junb* and *Il31ra* being differentially expressed amongst others in LCMV and MCMV-*ie2* infected mice 15 mpi, while *Ccl5, Uba52 and Ifi27l2a* were differentially expressed in both infections 28 dpi. This points towards potentially age-associated gene signatures with 100 and 52 differentially expressed genes shared between same-aged mice (Figure S4C), although this may be biased due to experimental batch effects. Next, we used the nearest-neighbor classifier of the ProjecTILs algorithm (Andreatta et al. 2021) to predict cell states within our data. ProjecTILs enables the projection of new scSeq data onto a previously generated and annotated reference T cell atlas and can therefore classify common T cell subsets (Andreatta et al. 2021). High proportions of predicted CD8+ effector memory cells could be detected in all mice, with the exception of 15 mpi with acute LCMV and CNC (Figure 3F, S5A). Moreover, the fraction of cells predicted to be CD8+ decreases at 15 mpi compared to their age-matched counterparts, again potentially representing the contraction occurring in virus specific T cells (Kaech, Wherry, and Ahmed 2002). Moreover, acute LCMV infections and CNC had high proportions of cells with a predicted CD8+ naive phenotype, which under experiment conditions most likely represents central memory cells (Andreatta et al. 2021). Both carrier-challenged mice showed higher proportions of T cells with a predicted exhaustion or exhausted precursor phenotype, even exceeding those arising from chronically infected mice 28 dpi (Figure 3F, S5A).

**Figure 3.**
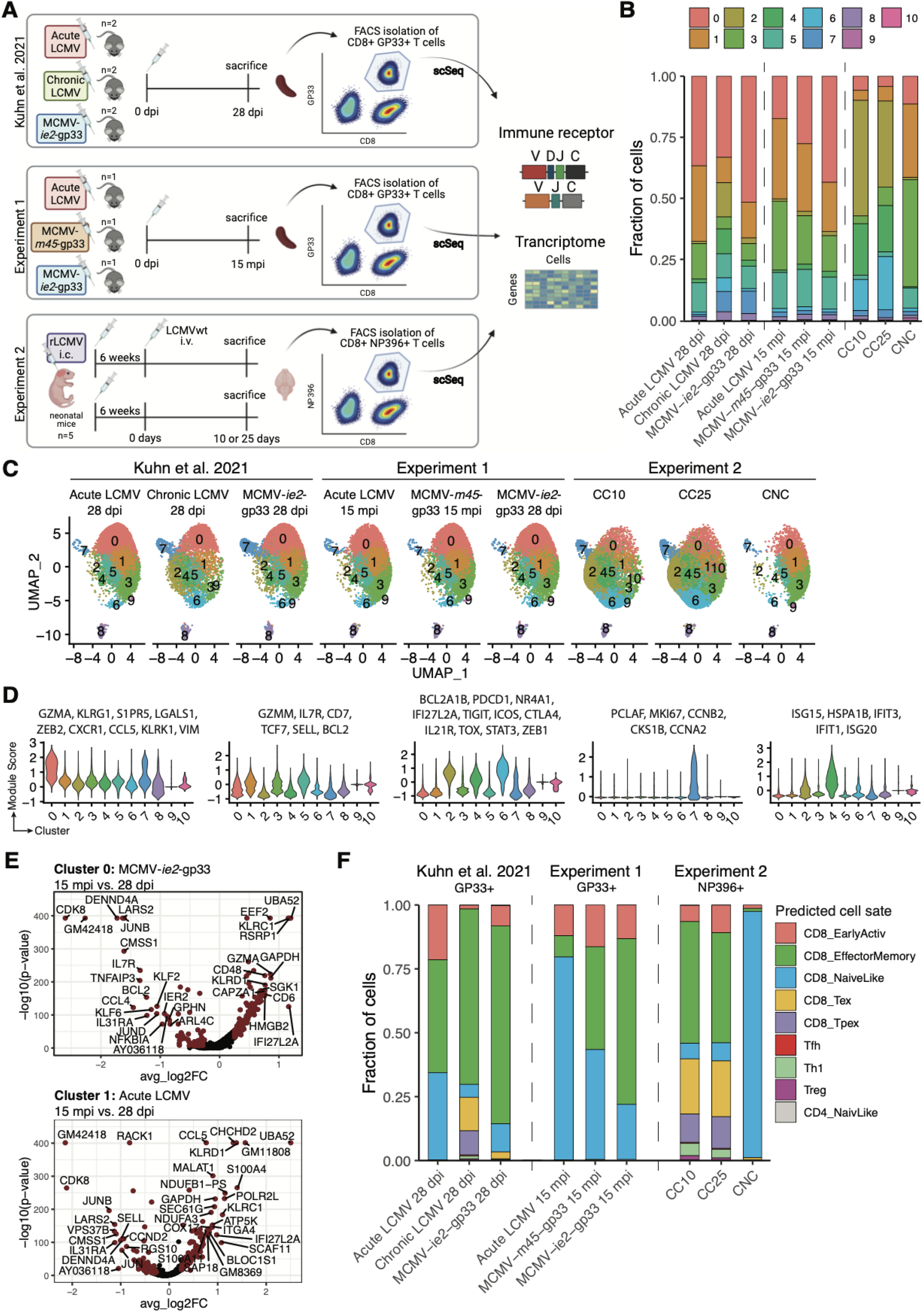
Virus-specific CD8+ T cells exhibit memory, inflationary, and exhausted phenotypes. **A**. Experimental Setup of each of the combined experiments. Previously published GP33-specific CD8+ cells from acute (n=2) and chronic (n=2) LCMV clone 13 and MCMV-*ie2*-gp33 (n=2) infected mice 28 days post infection (dpi) were integrated with GP33-specific CD8+ cells from acute LCMV clone 13 (n=1), MCMV-*m45*-gp33 (n=1), and MCMV-*ie2*-gp33 (n=1) infected aged mice (15 mpi). Additionally, NP396-specific CD8+ cells from carrier non-challenge (CNC) and carrier challenge 10 (n=2) and 25 (n=2) (CC10 and CC25, respectively) days post challenge from the viral déjà vu model. B. Fraction of cells belonging to each cluster in each infection type. C. Uniform manifold approximation projection (UMAP) split by infection type. D. Module-score for each cluster of genes associated with effector cell, memory, exhaustion, proliferative and interferon-expressing phenotypes. E. Differential gene expression within cluster 0 between MCMV-*ie2*-gp33 infection 28 dpi and 15 mpi (top). Differential gene expression within cluster 1 between acute LCMV infection 28 dpi and 15 mpi (bottom). Points in red indicate differentially expressed genes (adjusted p-value < 0.01 and average log_2_ fold change (FC) > 0.25). F. Fraction of cells belonging to predicted cell state using the nearest-neighbor classifier of the ProjecTILs algorithm in each infection type.

We were additionally interested in comparing sequence motifs of repertoires with various epitope-specificities and polyclonal repertoires. Comparison of the complementarity determining region 3 (CDR3) of TCR α and TCR β from GP33- and NP396-specific cells, as well as the previously discussed virus non-specific Tfh cells revealed biochemical differences between the groups (Figure S5B), again supporting that biochemical properties of TCR-interaction may dictate cell fate (Khatun et al. 2021).

### Clonally expanded T cells persist into old age and retain memory and inflationary phenotypes following acute and latent viral infections

After analyzing the global transcriptional landscape of our virus-specific CD8 T cell datasets, we questioned the extent of transcriptional heterogeneity within the different T cell populations persisting 15 mpi. Performing differential gene expression analysis between the aged groups emphasized the presence of the inflationary T cell phenotype (e.g., expression of *Klrg1, Gzma, Zeb2)* following infection with MCMV-*ie2-*gp33 compared to mice infected with acute LCMV or MCMV-*m45*-gp33. (Figure 4A). Moreover, non-inflationary T cells differentially expressed genes associated with effector phenotypes (*Gzmk, Ccl4*) compared to acute LCMV infection (Figure 4A). Having observed polyclonal but private clonal expansion in virus-specific cells 28 dpi, we questioned the extent to which clonally expanded cells were present in mice 15 mpi. Quantifying the fraction of the total repertoire composed by each clone demonstrated that highly expanded clones represented the vast majority of the repertoire for all three mice (Figure 4B). Interestingly, the most expanded clone in MCMV-*ie2*-gp33 infected mice represented ∼40% of the total recovered repertoire (Figure 4B). Analysis using ProjecTILs revealed a high proportion of CD8+ effector memory cells in MCMV infected mice compared to acute LCMV infection. Notably, MCMV-*m45*-gp33 infected mice had increased proportions of predicted naive-like cells (also representing central-memory cells) within the most expanded clones, again differentiating between inflationary and non-inflationary T cells (Figure 4B). We have previously reported that highly-expanded clones have relatively higher expression of effector molecules, such as *Nkg7, Ccl5*, and granzymes (Yermanos, Agrafiotis, et al. 2021; Yermanos, Neumeier, et al. 2021). This was consistent in MCMV-infected mice, however, was not observed at 15 mpi with acute LCMV infection.

**Figure 4.**
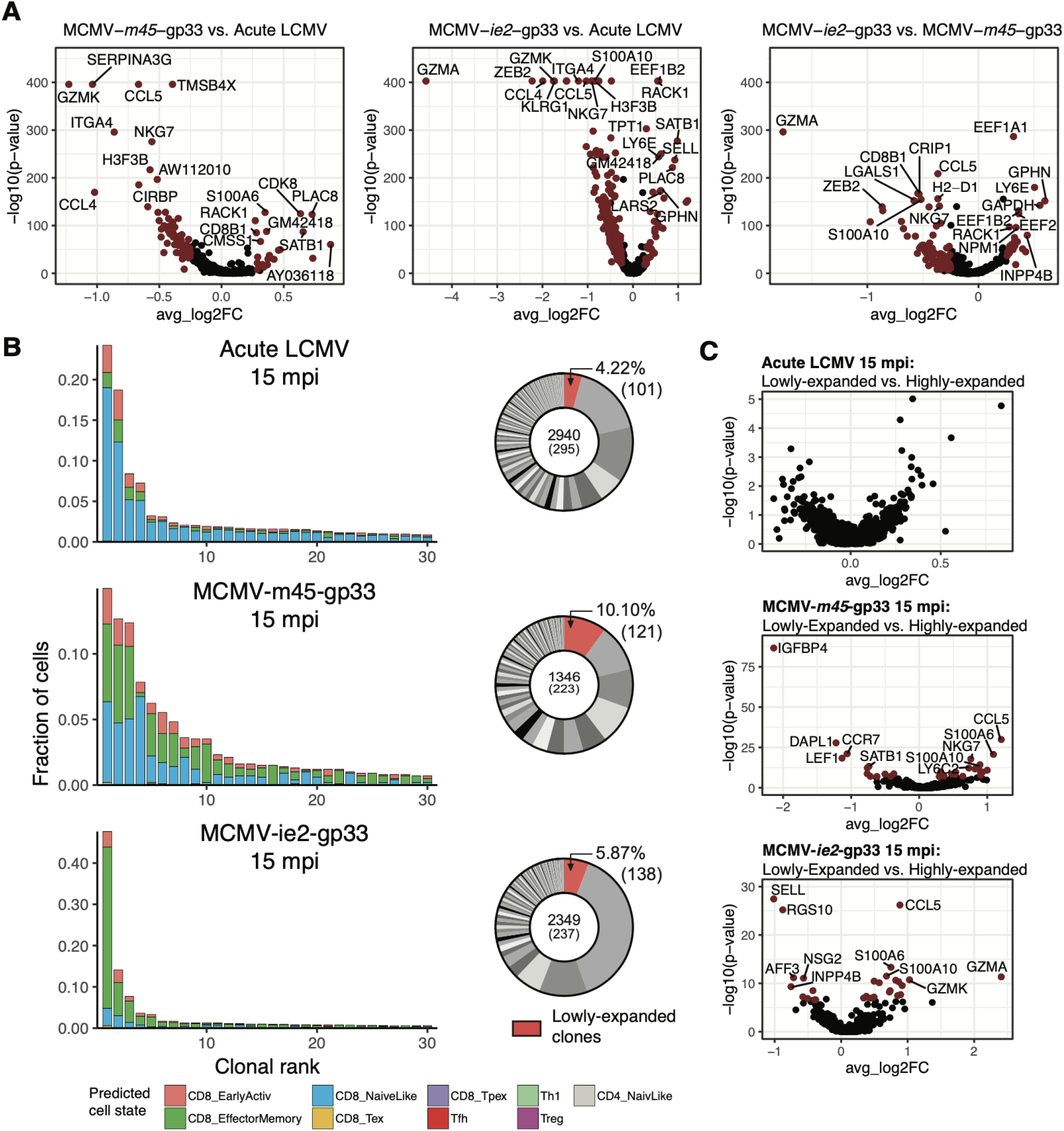
Memory and inflationary phenotypes in aged mice following acute and latent viral infections. A. Differential gene expression between aged mice. Points in red indicate differentially expressed genes (adjusted p-value < 0.01 and average log_2_ fold change (FC) > 0.25). B. Top 30 most expanded clones of aged mice separated by the predicted cell state using the nearest-neighbor classifier of the ProjecTILs algorithm (Andreatta et al. 2021). Distribution of clonal expansion is represented by donut plots. Each section corresponds to a unique clone (defined by CDR3α-CDR3β nt sequence) and the size corresponds to the fraction of cells relative to the total repertoire. Lowly-expanded clones (supported by only one unique cell) are colored in red. C. Differential gene expression within infection type between lowly- (=1 cell) and highly-expanded (>1 cell) clones.

### Carrier-challenged NP396-specific CD8+ cells retain an exhausted phenotype 25 days post viral challenge

In the viral déjà vu model, it has been reported that interferon-γ (IFN-γ) expression by CD8+ T cells and the resulting neuronal STAT1/CCL2 signaling play a key role in disease development (Kreutzfeldt et al. 2013; Di Liberto et al. 2018). However, the transcriptional programs and clonality of these virus-specific T cells have not been comprehensively profiled at a single-cell resolution. Performing differential gene expression analysis between all infection conditions revealed exhaustion and effector phenotypes (genes) in both CC10 and CC25 mice compared to CNC mice. Interestingly, both *Stat1* and *Ifng* were upregulated in CC25 compared to CC10 mice, despite earlier time points being associated with more severe clinical disease scores (Kreutzfeldt et al. 2013; Di Liberto et al. 2018). To explore clonal expansion during carrier-challenge, we quantified the fraction of the total repertoire composed by each clone. High levels of clonal expansion could be observed in all four mice. Within the top 30 most expanded clones, the majority of cells were predicted to have exhausted phenotypes by the ProjecTILs algorithm (Figure 5B). Moreover, even within the exhausted and effector memory cells, genes associated with TCR signaling (*Nra4a2, Nra4a3, Il21r, Stat1)* were differentially expressed 25 days post viral challenge.

**Figure 5.**
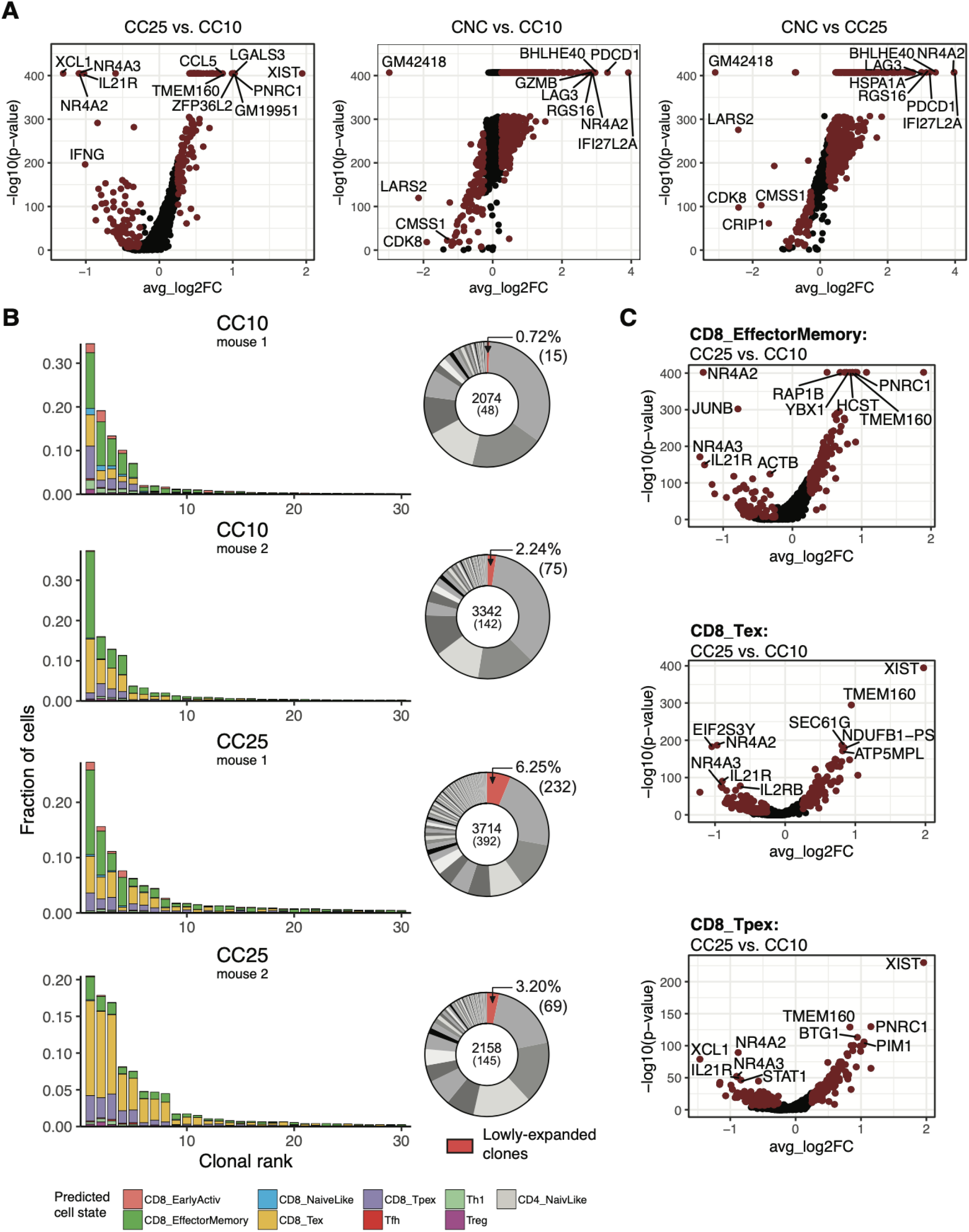
Carrier-challenged (CC) NP396-specific CD8+ exhibit exhausted cell signatures up to 25 days post viral challenge. A. Differential gene expression between mice from the viral déjà vu model. Points in red indicate differentially expressed genes (adjusted p-value < 0.01 and average log_2_ fold change (FC) > 0.25). B. Top 30 most expanded clones of mice from the viral déjà vu model separated by the predicted cell state using the nearest-neighbor classifier of the ProjecTILs algorithm (Andreatta et al. 2021). Distribution of clonal expansion is represented by donut plots. Each section corresponds to a unique clone (defined by CDR3α-CDR3β nt sequence) and the size corresponds to the fraction of cells relative to the total repertoire. Lowly-expanded clones (supported by only one unique cell) are colored in red. C. Differential gene expression within predicted cell states between CC25 and CC10.

### Lymphocytes are transcriptionally similar following EAE induction with either rMOG and MOG_33-55_

We finally investigated B and T cells during experimental autoimmune encephalomyelitis (EAE), a mouse model for the autoimmune neuroinflammatory disease multiple sclerosis (Constantinescu et al. 2011). In this model the autoreactive immune response can be elicited by immunization with a myelin-derived antigen coupled with an adjuvant. To this end, the recombinant myelin oligodendrocyte protein extracellular domain aa1-121 (rMOG) or the peptide MOG35-55 can be used amongst others. Notably, it has been suggested that B cells play differing roles in CNS pathology depending on whether rMOG or MOG35-55 is used (J. A. Lyons et al. 1999; J.-A. Lyons, Ramsbottom, and Cross 2002). Here, we performed single-cell sequencing of transcriptome and immune repertoires from CD3+ T and CD19+CD220+ B cells from the brains and spinal cords of mice immunized with rMOG + CFA and MOG35-55 + CFA, to compare how rMOG and MOG35-55 differentially imprint immune repertoires in the context of EAE (Figure 6A). To this end, we recovered GEX data for 5,444 T cells and 2,047 B cells with an average of 1,355 median genes per cell (Figure S6A). We first investigated common T cell markers and transcriptional cluster membership, which demonstrated similar expression patterns between rMOG and MOG35-55 groups (Figure 6B, 6C, S6B). Comparing cell states predicted using the ProjecTILs algorithm (Andreatta et al. 2021), similarly revealed high congruence between peptide- and protein-induced EAE (Figure 6D, S6C), with CD8+ effector memory and Th1 cells composing the majority of predicted cell states. Similar to the other T cell repertoires described within this study, we observed high levels of clonal expansion (Figure 6E). We further investigated whether gene signatures specific to expansion were present, which suggested that expanded CD4+ and CD8+ expanded T cells adopted a tissue-resident effector memory signature, associated with decreased expression of *Ilr7, Tcf7* and *S1pr1* and higher expression of *Cxcr6* (Figure 6S). A similar transcriptional signature was previously observed in expanded T cells in human MS patients (Beltrán et al. 2019). Expression of markers for mature B cells, such as *Cd19, Ebf1, Ms4a1* (encoding CD20) and *Ptprc*, was high in all clusters excluding clusters 5 and 6 (Figure 6F). After having observed transcriptional similarity across samples, we questioned whether similar patterns were also present in B cells. Cells from both immunizations were present in all resulting transcriptional clusters (Figure 6E). The clonal expansion of B cells expressing a single heavy and light chain was lower compared to T cells, while expansion between immunization strategies did not differ (Figure 6H). Assessing the isotypes of the top 30 most expanded B cells clones, in addition to all clones, revealed high frequencies of the IgM isotype (Figure 6G, 6H), which is similar to previous findings profiling B cells in the CNS of naive mice (Yermanos, Neumeier, et al. 2021).

**Figure 6.**
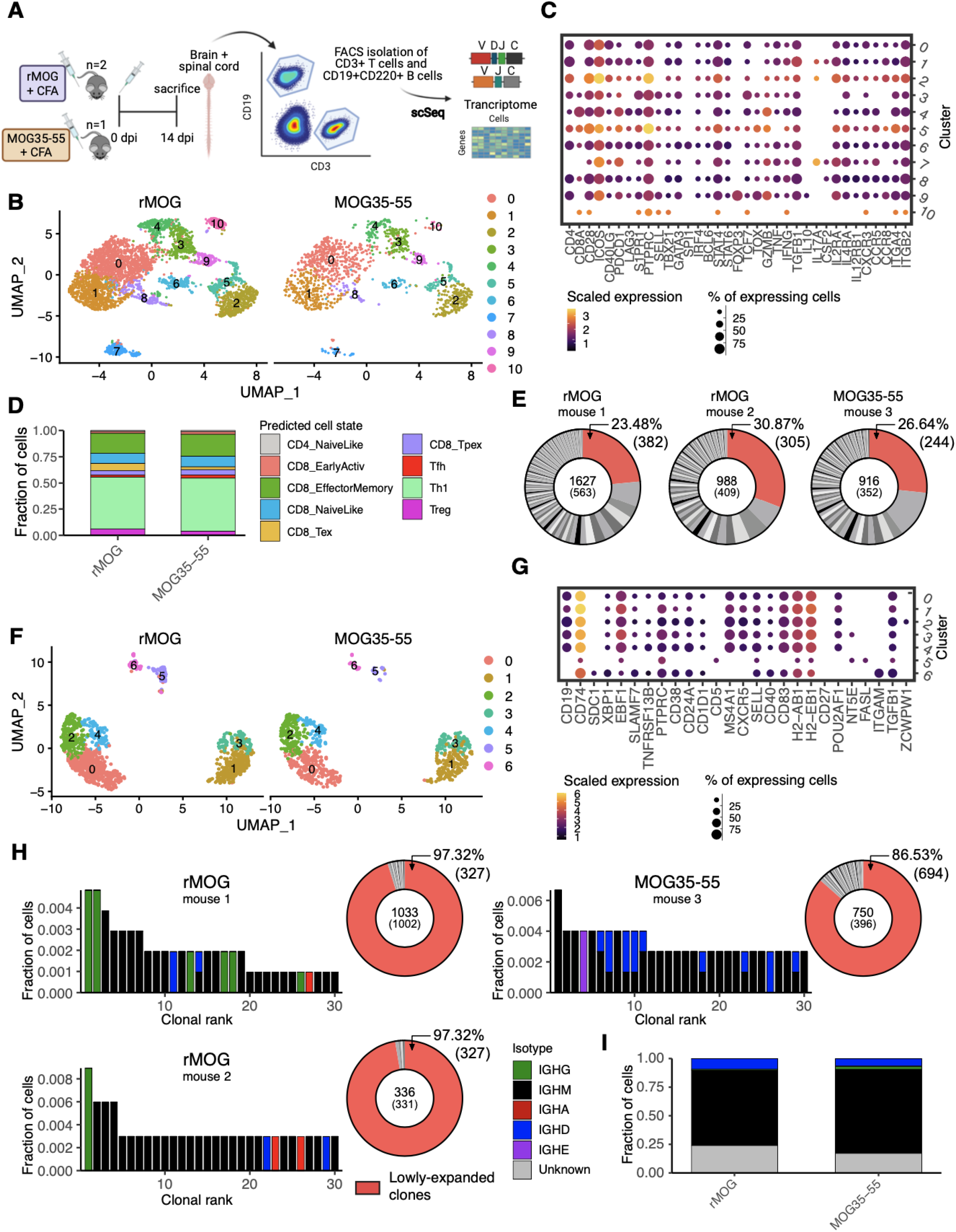
Transcriptional similarity of B and T cells following experimental autoimmune encephalomyelitis (EAE) induction with either rMOG and MOG_33-55_. A. Experimental setup of single-cell immune repertoire sequencing. CD3+ T and CD19+CD220+ B cells were isolated from mice immunized with either recombinant myelin oligodendrocyte protein extracellular domain aa1-121 (rMOG, n=2) or the peptide MOG35-55 (n=1), after 14 days.B. Uniform manifold approximation projection (UMAP) of T cells split by immunization type. C. Dottile plot showing genes of particular interest in all clusters arising from unsupervised clustering in T cells. D. Fraction of T cells belonging to predicted cell state using the nearest-neighbor classifier of the ProjecTILs algorithm (Andreatta et al. 2021) for both groups. E. Donut plots representing clonal expansion of T cells. Each section corresponds to a unique clone (defined by CDR3α-CDR3β nt sequence) and the size corresponds to the fraction of cells relative to the total repertoire. Lowly-expanded clones (supported by only one unique cell) are colored in red. F. Uniform manifold approximation projection (UMAP) of B cells split by experimental group G. Dottile plot showing genes of particular interest in all clusters arising from unsupervised clustering in B cells. H. Top 30 most expanded B cell clones of mice immunized with either rMOG or MOG35-55 colored by isotype. Distribution of clonal expansion is represented by donut plots. Each section corresponds to a unique clone (defined by CDRH3-CDRH3 nt sequence) and the size corresponds to the fraction of cells relative to the total repertoire. Lowly-expanded clones (supported by only one unique cell) are colored in red. I. Fraction of B cells corresponding to a specific isotype. Cells with either less or more than one heavy and one light chain were assigned an “unknown” isotype.

## Discussion

Single-cell sequencing of lymphocytes now makes it possible to profile adaptive immune repertoires and their transcriptomes at high-throughput. Immune repertoire features such as clonal expansion, germline gene usage, and transcriptional phenotypes provide a quantitative overview of an immune response and hold the potential to infer immunological status. Despite a growing number of studies leveraging single-cell immune repertoire sequencing, some pilot and small-scale sequencing experiments do not reach the public domain despite the potential to both uncover immunological principles and integrate with larger public datasets. Although low numbers of mice limit the conclusions that can be made by this study, our single-cell sequencing data presented can nevertheless be used in combination with other immune repertoire data as demonstrated by multiple of our conclusions aligning with previously published results. Examples of this include transcriptional differences between early and later time points of Tfh (Crotty 2014), GP66-specific TCRs detected in Tfh following LCMV infection (Khatun et al. 2021), distinct repertoire features of Tfr (Maceiras et al. 2017), inflationary and exhaustion phenotypes of virus-specific CD8+ T cells (Welten et al. 2020; Merkler et al. 2006), and minor differences between B and T cells when inducing EAE with either rMOG or MOG35-55 (Sefia et al. 2017). In addition to these confirmatory findings, our collection of single-cell immune repertoire sequencing can spark future experiments investigating clonal selection of lymphocytes across various experimental models. These include relating antigen specificity to expansion and expression phenotypes of Tfh populations during chronic infection, quantifying repertoire dynamics relating to Tfh and Tfr during germinal center collapse, exploring consequences of prolonged T cell exhaustion and effector phenotypes persisting beyond the peaks of neurological disease (déjà vu) and latent viral infections, or revisiting previously reported differences between MOG peptide and protein models of EAE in terms of the adaptive immune response (J. A. Lyons et al. 1999).

## Methods

### Mice Experiments

All animal experiments were performed in accordance with institutional guidelines and Swiss federal regulations. Experiments involving LCMV, MCMV-*ie2*-gp33, MCMV-*m45*-gp33 infections were approved by the veterinary office of the canton of Zurich or Geneva under animal experimentation licenses ZH114/2017, ZH115/2017 and GE170-19. Female C57BL6J WT mice between the age of 8-12 weeks were used for all experiments unless otherwise stated. Acute LCMV infections were infected intravenously (i.v.) with 200 focus forming units (ffu) of LCMV clone 13 in the tail vein. Latent infections were established by injecting 2×10^5^ pfu dose of MCMV-*ie2*-gp33 or MCMV-*m45*-gp33 i.v., which were obtained from Dr. L. Cicin-Sain and contained a functional *m157* gene as previously described (Welten et al. 2015). MCMV viral stocks were propagated on M2-10B4 cells and purified by ultracentrifugation using a 15% sucrose gradient. LCMV clone 13 was produced as previously described (Battegay et al. 1993; Sandu et al. 2020). Upon sacrifice 28 dpi or 15 mpi, organs were harvested and single-cell suspensions were prepared by mashing the tissue through a 70 µM cell strainer and rinsing with complete RPMI (RPMI-1640 supplemented with 10% fetal bovine serum, 2 mM L-glutamine, 1% penicillin-streptomycin, 1 mM sodium pyruvate, 50 nM beta-mercapthoethanol, 0.1 mM non-essential [glycine, L-alanine, L-asparagine, L-aspartic acid, L-glutamic acid, L-proline, L-serine] amino acids, 20 mM HEPES). The single-cell suspension was then incubated with CD8-PE (clone 53-6.7, Biolegend), MHC class 1 tetramer for gp_33-41_ conjugated to APC diluted in FACS buffer (PBS, 2 mmEDTA, 2% FCS) at room temperature for 30 minutes, as previously described (Altman et al. 1996), and LiveDead nearIR (Thermo Fisher). Tetramer-positive cells were isolated via flow cytometric sorting (FACSAria with FACSDiva software) and subsequently supplied as input for single-cell immune repertoire sequencing. Recombinant LCMV strains and viral déjà vu infections were set up as previously described (Di Liberto et al. 2018). EAE was induced by the injection of either 325 μg rMOG or 200 μg MOG35-55 in CFA as previously described (Knier et al. 2018). Following sacrifice, mice were transcardially perfused with 4°C cold PBS for 3 minutes. Brains and spinal cords were collected in RPMI, transferred to a tube containing 1mL RPMI with 1.0 mg/ml Collagenase I/DNAse I, cut into small pieces, incubated for a further 30 min at 37 °C and mashed through a 70 μm cell strainer. Single-cell suspensions of brains and spinal cords from each mouse were pooled in the case of rMOG. Lymphocytes from brain and spinal cord for déjà vu and EAE models were isolated using 30% and 70% Percoll gradients by centrifugation. Cells were stained with anti-CD3-FITC, anti-B220-APC and anti-CD19-PE-Cy7 and sorted on a FACSAria 3 (BD Biosciences) as B cells (CD19+, B220+) or T cells (CD3+, CD19-) into 1.5ml tubes.

### Single-cell immune repertoire analysis

Single-cell immune repertoire sequencing was performed as according to the 10x Genomics Chromium Single Cell V(D)J Reagents Kit (CG000166 Rev A) as previously described (Neumeier et al. 2021). In brief, single cells for all samples were simultaneously encapsulated with gel emulsion microdroplets (10x Genomics, 1000006) in droplets using a Chromium Single Cell A Chip (10x Genomics, 1000009) with a target loading of 13,000 cells per reaction. cDNA amplification was performed using 14 cycles and subsequently split for downstream GEX and VDJ library preparation. GEX libraries were amplified using the Chromium Single Cell 5’ Library Kit (10x Genomics, 1000006). TCR libraries were amplified using the Chromium Single Cell V(D)J Enrichment Kit, Mouse T Cell (10x Genomics, 1000071). Final libraries were pooled and sequenced on the Illumina NovaSeq S1 using a concentration of 1.8 pM with 5% PhiX. Paired-end sequencing files for GEX and VDJ libraries were aligned to the murine reference genome (mm10) and V(D)J germlines (GRCm38) using 10x Genomics cellranger (v5.0.0) count and vdj arguments, respectively. The filtered feature matrix directory was supplied as input to the VDJ_GEX_matrix function in the R package Platypus (v3.2.1) (Yermanos, Agrafiotis, et al. 2021), which uses the transcriptome analysis workflow of the R package Seurat (Satija et al. 2015). Only those cells containing less than 20% of mitochondrial reads were retained in the analysis. Genes involved in the adaptive immune receptor (e.g., TRB, TRBV1-1), were removed from the count matrix to prevent clonal relationships from influencing transcriptional phenotypes. Gene expression was normalized using the “scale.data” (Tfh, EAE) or “harmony” (GP33+ aged, déjàvu) argument in the VDJ_GEX_matrix function. 2000 variable features were selected using the “vst” selection method and used as input to principal component analysis (PCA) using the first 10 dimensions. Graph-based clustering using the Louvain modularity optimization and hierarchical clustering was performed using the functions FindNeighbors and FindClusters in Seurat using the first ten dimensions and a cluster resolution of 0.5. UMAP was similarly inferred using the first ten dimensions. The FindMarkers function from Seurat was used when calculating differentially expressed genes (both across groups or across clusters) with logfc.threshold set to 0 and minimum number of cells expressing each gene set to 0.25 and subsequently supplied to the GEX_volcano and GEX_gsea functions from Platypus. Mitochondrial and ribosomal genes were removed when visualizing DE genes. Gene expression heatmaps, feature plots, and violin plots were produced by supplying genes of interest to the functions DoHeatmap, FeaturePlot, and VlnPlot functions in Seurat, respectively. Module score was calculated using the AddModuleScore from Seurat. The GEX_gsea function uses the R package fgsea (v1.16.0) (Sergushichev 2016), which performs gene set enrichment analysis (GSEA) and uses the adaptive multilevel splitting Monte Carlo approach. Hallmark (H) gene sets from the Molecular Signatures Database (MSigDB) were used for GSEA (Subramanian et al. 2005). Cells containing no or more than one α/heavy and β/light chain were filtered out for TCR/BCR repertoire analysis. Clones were defined by identical CDR3α/CDRH3 and CDR3β/CDRL3 sequence (nucleotide or amino acid sequence) across all repertoires. Clones represented by more than one cell were considered highly-expanded clones, while single-celled clones were defined as lowly-expanded. The VDJ_circos function from Platypus was used to create circos plots. The R package pheatmap (v1.0.12) was used to create heatmaps displaying V gene usage or number of shared clones (Kolde and Kolde 2015). The projection of cells onto reference UMAPs and cell state predictions were done using the R package ProjecTils (Andreatta et al. 2021), Each experiment was projected individually onto the tumor-infiltrating T lymphocytes (TIL) atlas.

## Data visualization

Donut plots were created using GraphPad Prism® Software version 9. The packages ggplot (v3.3.3), ggrepel (v0.9.1), ggpubr (v0.4.0), cowplot (v1.1.1) and gridExtra (v2.3), were used for data visualization. Experimental overviews were created with BioRender.com.

## Data Availability

All GEX and VDJ immune repertoire sequencing data has been deposited on the EMBL’s European Bioinformatics Institute website under the accession code E-MTAB-11330.

## Acknowledgements

We acknowledge and thank Dr. Christian Beisel, Elodie Burcklen, Ina Nissen, and Mirjam Feldkamp at the ETH Zurich D-BSSE Genomics Facility Basel for excellent support and assistance. We also thank Nathalie Oetiker, Franziska Wagen, Gregory Schneiter, and Anne Häfke for excellent experimental support. Further we acknowledge Prof. Martin Kerschensteiner for providing rMOG protein used for EAE experiments.

## Author contributions

DS, KH, AA, VK, MMC, RD, RK, IS, GS, NP, NF, MP, IV, AM, BK, IW, RE, LF, KK, DN, and AY performed experiments and/or computational analyses. All authors contributed to designing the study and writing the manuscript.

---

**Figure S1.**
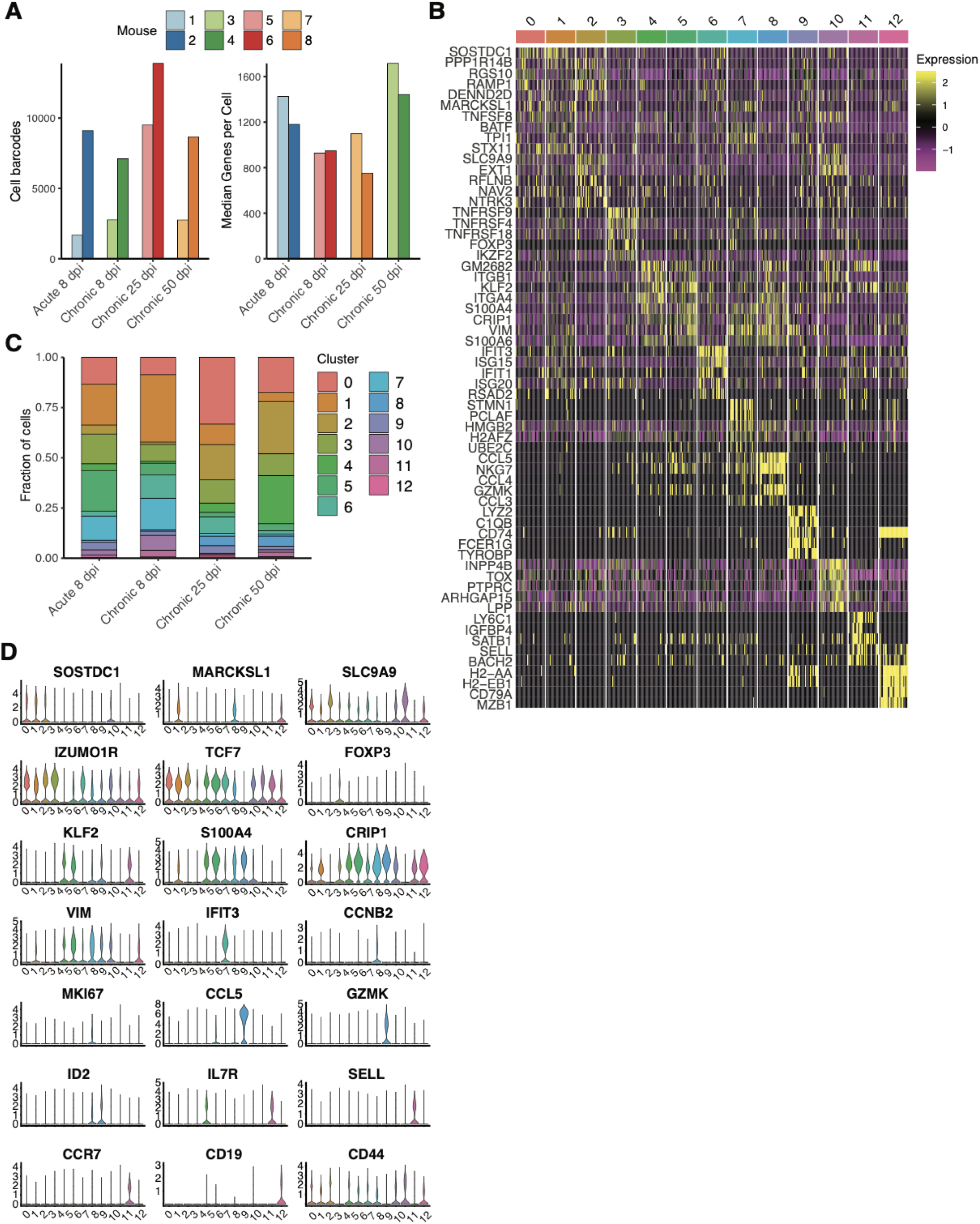
Cluster defining gene signatures in Tfh cells after acute and chronic infection. A. Number of cell barcodes with gene expression information for each mouse (left) and median number of genes per cell for each mouse (right). B. Fraction of cells belonging to each cluster for each experimental group. C. Top five significant genes defining each cluster ranked by average log fold change. D. Normalized expression for genes of particular interest in each cluster.

**Figure S2.**
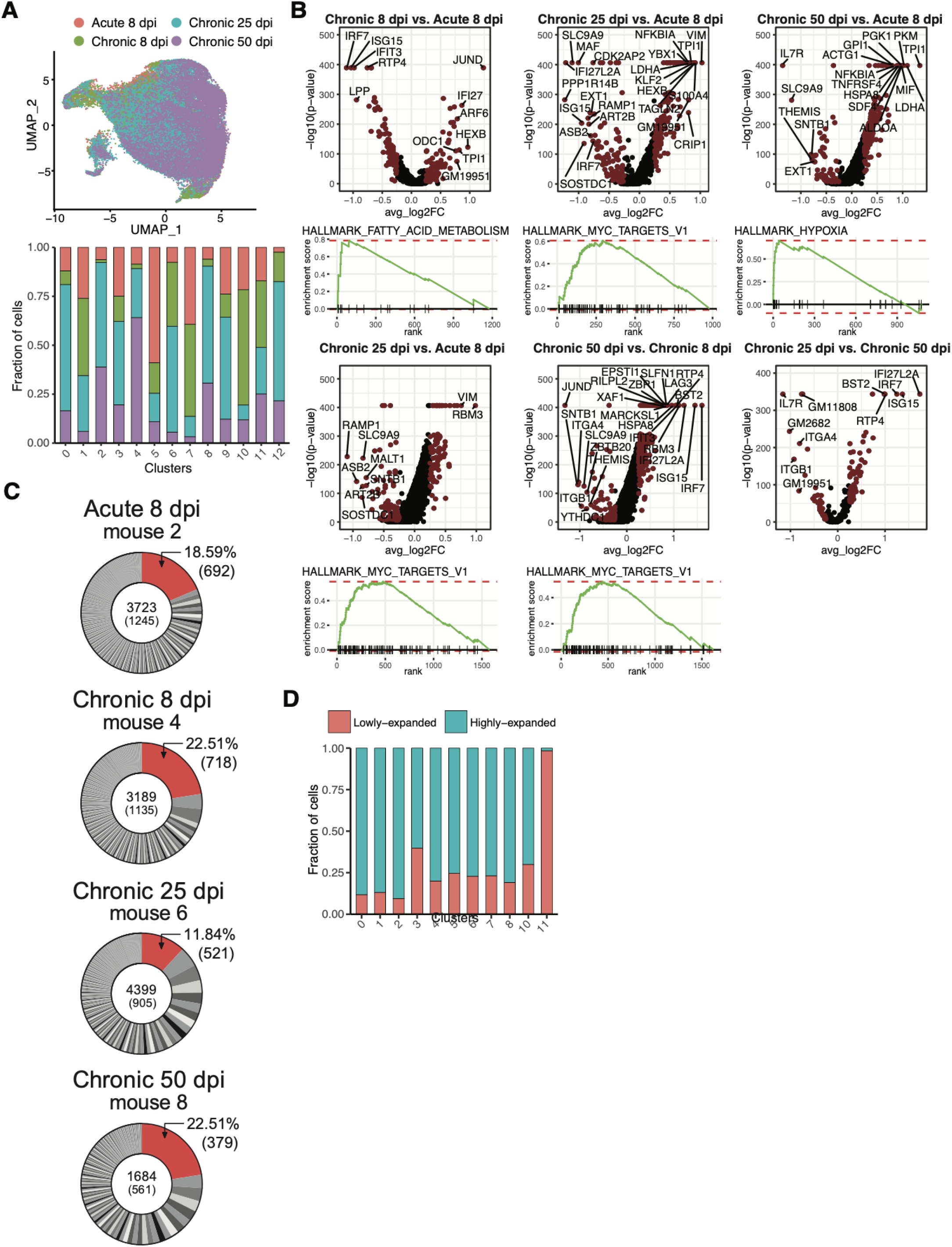
Differential gene expression of Tfh cells during early and late LCMV infection. **A**. Uniform manifold approximation projection (UMAP) displaying infection time point (top) and fraction of cells belonging to each infection time point in each transcriptional cluster. B. Differential gene expression and gene set enrichment (GSEA) analysis between acute LCMV 8 dpi and chronic LCMV 8 dpi (top left), acute LCMV 8 dpi and chronic LCMV 25 dpi (top middle), acute LCMV 8 dpi and chronic LCMV 50 dpi (top right), chronic LCMV 8 dpi and chronic LCMV 25 dpi (bottom left), chronic LCMV 8 dpi and chronic LCMV 50 dpi (bottom middle) and chronic LCMV 25 dpi and chronic LCMV 50 dpi (bottom right). Points in red indicate differentially expressed genes (adjusted p-value < 0.01 and average log_2_ fold change (FC) > 0.25). C. Residual donut plots showing distribution of clonal expansion in each infection time point. Each section corresponds to a unique clone (defined by CDR3α-CDR3β nt sequence) and the size corresponds to the fraction of cells relative to the total repertoire. Lowly-expanded clones (supported by only one unique cell) are colored in red. D. Fraction of lowly- and highly-expanded cells in each transcriptional cluster.

**Figure S3.**
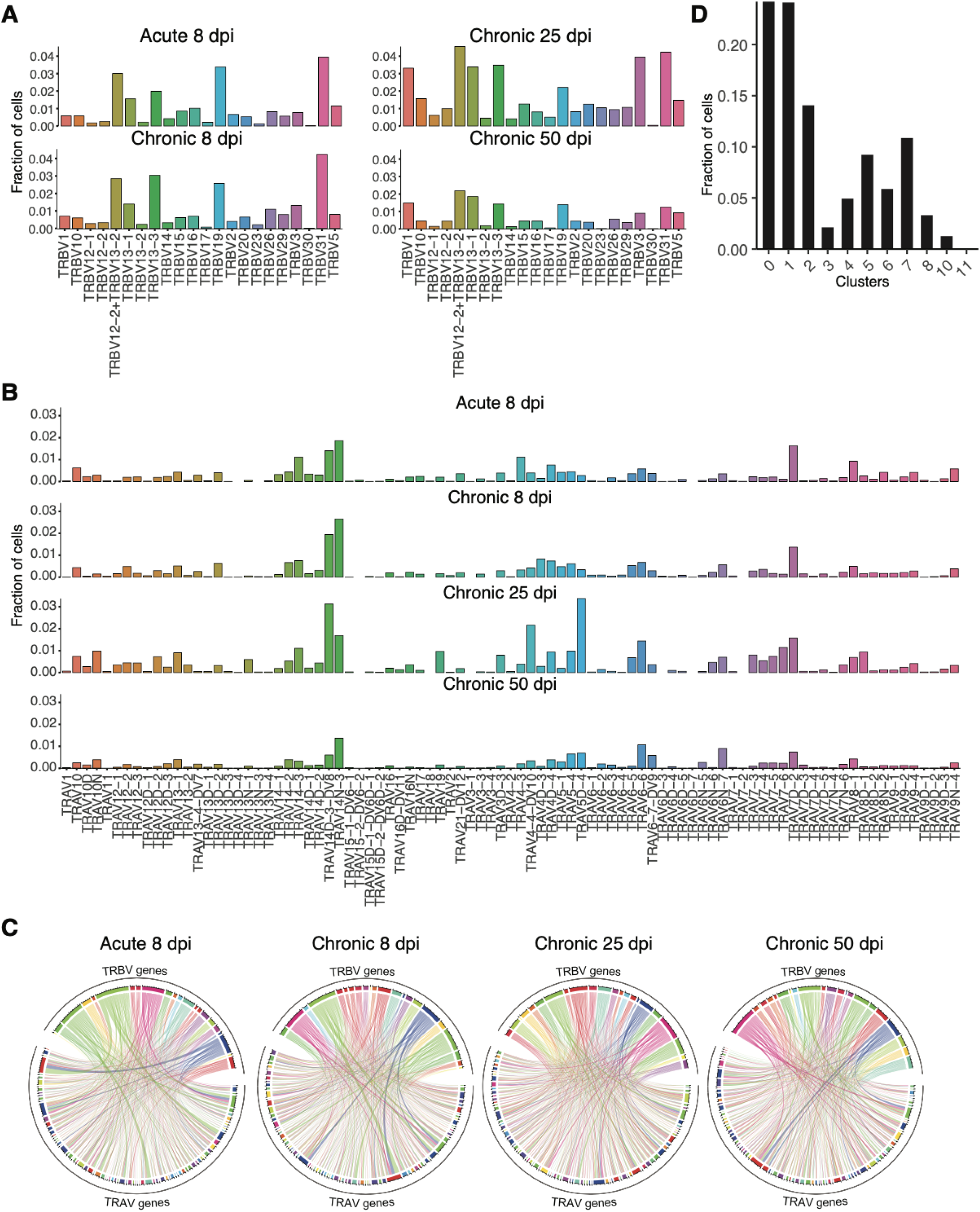
Tfh cells during acute and chronic LCMV infection show similar abundance of TRAV and TRBV genes but no dominant pairing patterns. A. TRBV germline gene usage within Tfh clones (CDR3α-CDR3β nt sequence). B. TRAV germline gene usage within Tfh clones (CDR3α-CDR3β nt sequence). C. Circos plots depicting the relationship between TRB and TRA V genes. Color corresponds to TRBV gene usage. Connections illustrate the number of cells using each particular combination. D. Fraction of shared clones within each transcriptional cluster.

**Figure S4.**
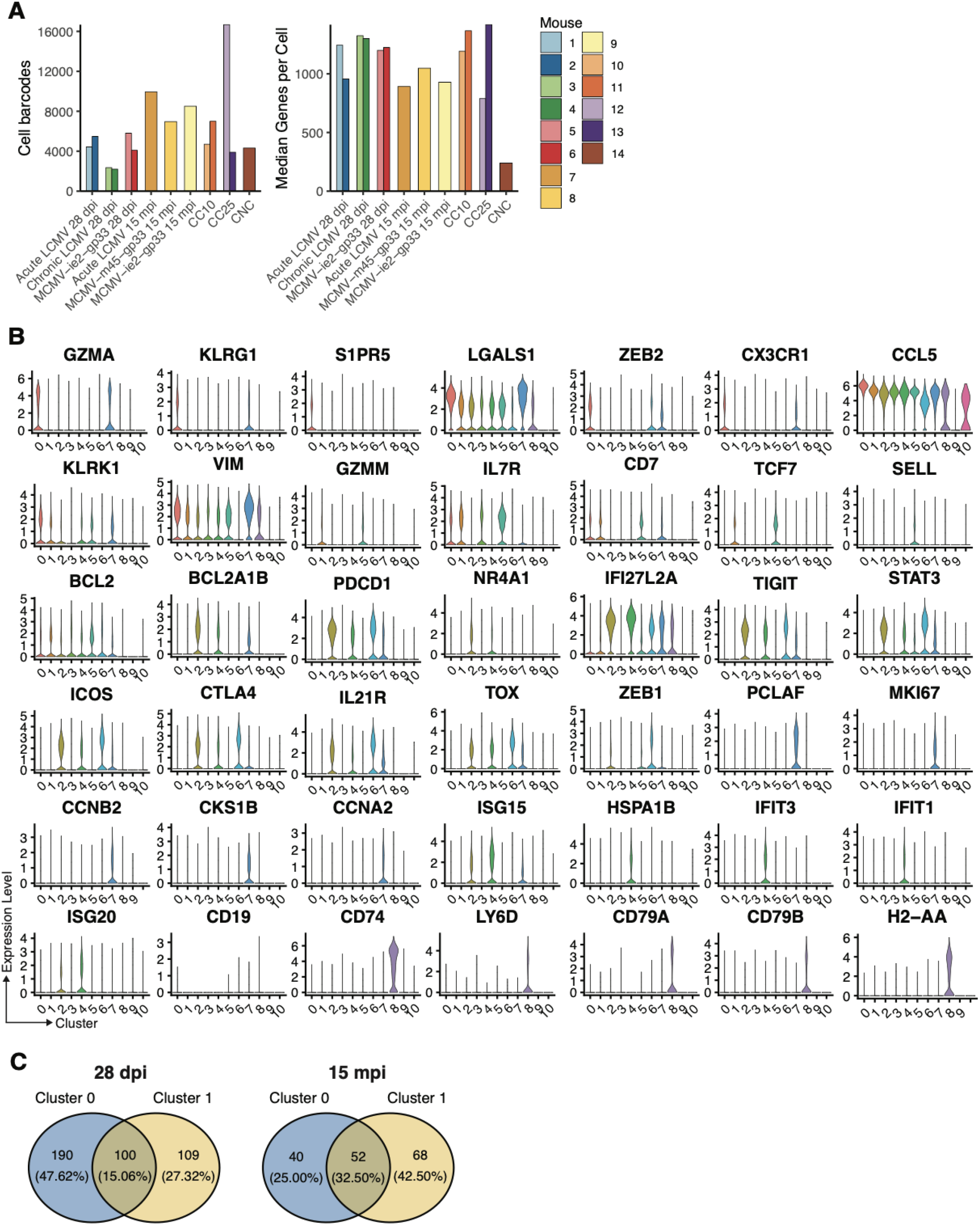
Cluster defining gene signatures in virus-specific CD8+ T cells. **A**. Number of cell barcodes with gene expression information for each mouse (left) and median number of genes per cell for each mouse (right). B. Normalized expression for genes of particular interest in each cluster. C. Venn diagram depicting number and percentage of differentially expressed genes shared between clusters of same-aged mice.

**Figure S5.**
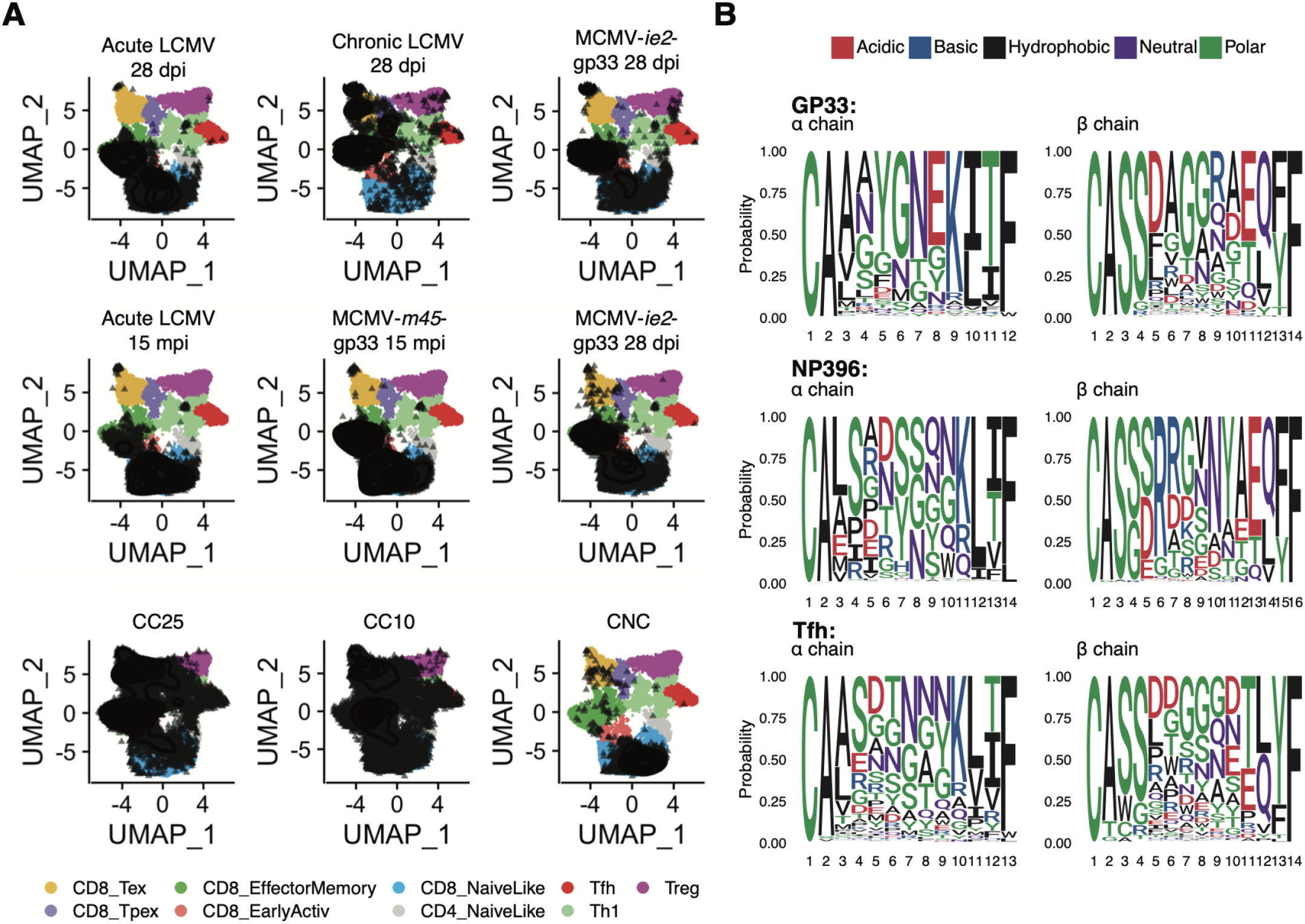
Predicted Cell states and repertoire sequence motifs of virus-specific CD8+T cells and polyclonal Tfh cells. A. Single-cell projection of GP33- and NP396-specific CD8+ T cells on the default reference TIL atlas (Andreatta et al. 2021). B. Sequence logo plots of the alpha and beta chain of GP33- and NP396-specific CD8+ T cells and polyclonal Tfh cells.

**Figure S6.**
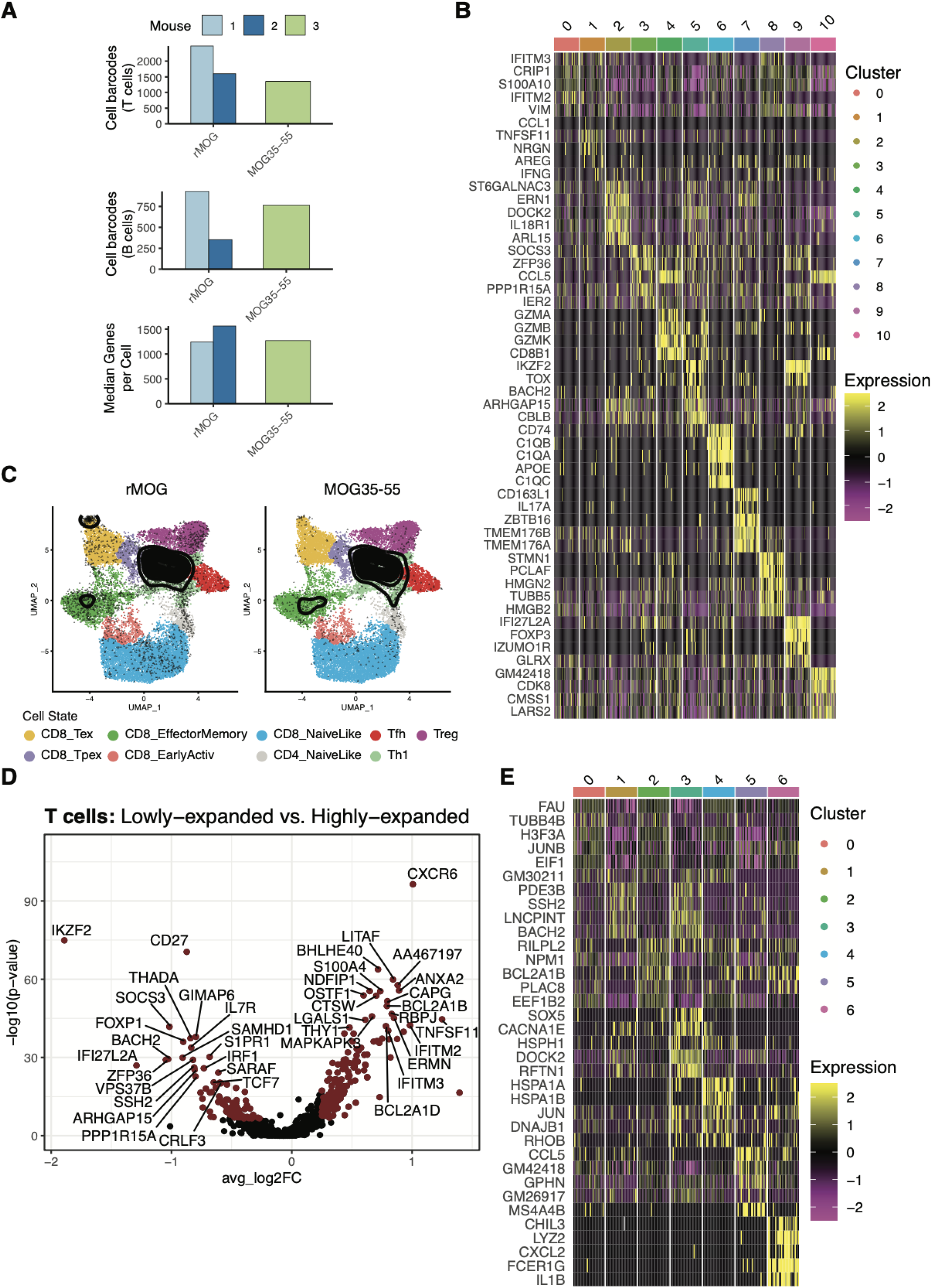
Cluster defining gene signatures in B and T cells following experimental autoimmune encephalomyelitis (EAE) induction with either rMOG and MOG_33-55_. A. Number of cell barcodes with gene expression information for each mouse and median number of genes per cell for each mouse. B. Single-cell projection of T cells following EAE induction with either rMOG and MOG_33-55_ on the default reference TIL atlas (Andreatta et al. 2021). C. Top five significant genes defining each cluster of T cells ranked by average log fold change. D. Differential gene expression between lowly-expanded (1 cell) and highly-expanded (more than one cell) CD4+ and CD8+ T cells E. Top five significant genes defining each cluster of B cells ranked by average log fold change

